# Protein-protein interaction (PPI) network analysis reveals important hub proteins and sub-network modules for root development in rice (*Oryza sativa*)

**DOI:** 10.1101/2022.06.06.494990

**Authors:** Samadhi Wimalagunasekara, Shamala Tirimanne, Pasan Chinthana Fernando

## Abstract

The root system is vital to plant growth and survival. Therefore, genetic improvement of the root system is beneficial for developing stress-tolerant and improved plant varieties. This requires the identification of proteins that significantly contributes to root development. Analyzing protein-protein interaction (PPI) networks is vastly beneficial in studying developmental phenotypes, such as root development because a phenotype is an outcome of several interacting proteins. PPI networks can be analyzed to identify modules and get a global understanding of important proteins governing the phenotypes. PPI network analysis for root development in rice has not been performed before and has the potential to yield new findings to improve stress tolerance. Therefore, in this research, the network module for the root development was extracted from a PPI network retrieved from the STRING database. Novel protein candidates were predicted, and hub proteins and sub-modules were identified from the extracted module. The validation of the predictions yielded 75 novel candidate proteins, 6 sub-modules, 20 intramodular hubs, and 2 intermodular hubs. These results show how the PPI network module is organized for root development and can be used for future wet-lab studies for producing improved rice varieties.

## 1. Introduction

*Oryza sativa* has a very high demand as a staple food, and although much successful research has been carried out to improve the yield, it tends to decrease drastically in response to environmental stresses. Therefore, improvement of *O. sativa* must continue, aiming at sufficient supply for the increasing population, which requires producing improved *O. sativa* varieties with higher yields and a higher ability to withstand environmental stresses [1–3].

The root system is the main component that supplies water and nutrients to the plant and plays a major role in withstanding environmental stresses [4]. Therefore, the root system should have a high priority when improving plant varieties. Root development is a complex biological process regulated by a collection of biological pathways, which are influenced by environmental and genetic factors [4]. This research is focused on investigating the genetic factors by identifying the functionally important proteins and their interactions responsible for root development in *O. sativa* using network analysis.

Biological processes are regulated by a collection of proteins and their interactions. These protein-protein interactions (PPI) are represented as PPI networks [5,6]. PPI data are generated using wet-lab and computational techniques and are stored in databases [7,8], such as DIP, STRING, and BioGRID. Among these databases, the STRING database is popular because of the higher abundance, coverage, and better quality control of PPI data [7,9,10]. STRING contains PPIs from both experimental methods and computational methods and provides a quality score for each interaction by integrating the data from various resources such as literature and gene expression profiles [7,10,11]. PPI networks contain modules, which are distinct collections of proteins usually specific for a particular function or a phenotype [5,12]. Hence, PPI networks can be analyzed to identify modules, which represent underlying protein interactions that determine the molecular functions and phenotypes. Furthermore, PPI networks are used for predicting novel protein candidates for molecular functions and phenotypes based on their interactions with known neighbors [13,14]. Though wet lab methods are available for predicting new protein candidates, computational methods for protein function prediction are faster, more cost-effective, and less laborious than wet lab methods [15,16]. Sequence similarity-based methods are popular computational approaches, which have been proven to be effective for some protein molecular function prediction studies, but they are less efficient for phenotype studies [15,17]. This is because proteins associated with one phenotype can include proteins with highly diverse sequences, annotated with different molecular functions [14,17]. Therefore, predicting proteins for phenotypes using PPI networks is more accurate and comprehensive than sequence similarity-based methods [15,17].

PPI network analysis can be used to identify the sub-modules within a particular module of a phenotype, and analysis of these sub-modules allows to identify the related biological pathways and important proteins, i.e., hub proteins, involved in those pathways [18,19]. Identifying hub proteins of a module is an important advantage of performing network module analysis [14]. Hubs have a higher number of interactions compared to non-hubs [20]. There are two types of hubs: intramodular and intermodular hubs [21]. Intramodular hubs are can be found with their partners within the functional modules while intermodular hubs act among the modules and interconnect them [20–22]. Removal of a hub has a higher impact compared to non-hubs because it impacts several biological pathways in the network [21], which disrupts the resulting phenotype. Therefore, hub proteins are identified as important proteins that play a critical role in maintaining module organization and stability [23]. These are usually important drug targets in human-related studies [24] or genetic engineering targets in crop improvement [25].

PPI networks allow the understanding of the global organization of PPIs, sub-modules, connectivity among those sub-modules, and the hub proteins [8,26,27]. The interpretation of these networks reveals the biological pathways associated with a particular phenotype. The efficiency of PPI network analysis has been proven in human disease research [8,28,29], but to our knowledge, this method has never been used on root development in *O. sativa*. Therefore, this study was conducted to understand the PPI network module of root development in *O. sativa*.

The analysis included predicting protein candidates for root development, extracting, and visualizing the module, identifying sub-modules, analyzing their biological pathways, and identifying important hub proteins and their interaction pathways for root development in *O. sativa*. This large-scale analysis reveals how PPIs are responsible for root development, which can be used for future studies to develop new and improved stress-resistant *O. sativa* varieties.

## 2. Materials and methods

### 2.1. Data retrieval and pre-processing

The proteins already known to be involved in root development (seed proteins) were retrieved from the literature and the STRING database (version 11.0; July 2019; https://string-db.org/). The PPI network and supplementary data for *O. sativa* were downloaded from the STRING database. (Retrieved and downloaded on July 17, 2019)

To improve the reliability and the quality of the downloaded PPI network, it was filtered using the ‘combined score’ according to a standard cutoff mark of 400 [10]. Duplicates of the same record were removed and STRING identifiers (IDs) for proteins were converted to preferred protein names to facilitate further analysis.

### 2.2. Network-based candidate gene prediction and root development protein module extraction

Hishigaki method was selected for the candidate gene prediction [13] and the prediction score was calculated according to the equation below.

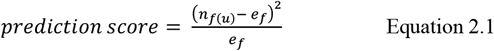

In Equation 2.1, *f* denotes the function of interest, and *u* denotes the protein of interest. The number of proteins with the function (*f*) in the n-neighborhood of *u* is given by *n_f(u)_* and *e_f_* denotes the expected frequency for the function calculated as follows:

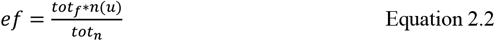

In Equation 2.2, the total number of proteins annotated to the function of interest (*f*) in the network is denoted by *tot_f_. Tot_n_* denotes the total number of proteins in the network; *n(u)* denotes the total number of proteins in the immediate neighborhood of the interested protein (*u*) [13].

Predicted scores were sorted to obtain the proteins with the highest predicted scores. Top 20, 50, 75, and top 100 proteins were listed with seed proteins and PPI modules for those lists were extracted from the pre-processed network and visualized using the Cytoscape software (version 3.7.1) [30,31]. The top protein cutoff with better network connections and degree distributions was selected for the final analysis based on the visualizations.

### 2.3. Validation of the predictions

Computational validation of the predicted proteins was required to confirm the accuracy of the predictions. Validation was done using enrichment analysis and performing a literature search on the predicted proteins.

Enrichment analyses were performed using the DAVID (DAVID Bioinformatics Resources 6.8; https://david.ncifcrf.gov/home.jsp) web application. The functional annotation tool in DAVID was used and the official gene symbol was selected as the identifier [32]. The biological process component of the Gene Ontology (GO-BP) and KEGG pathways, which had significant p-values (>0.05), were selected for further analysis [32]. Literature searches were also used to further validate the predictions and the enriched biological pathways.

Furthermore, receiver operating characteristic (ROC) curves and precision-recall curves were generated to evaluate the performance of the Hishigaki prediction method using leave-one-out cross-validation and seed proteins [33,34].

### 2.4. Identification and analysis of sub-modules

Preliminary identification of sub-modules was done using MCODE (version 1.5.1) [35,36] plug-in in Cytoscape software with the clustering parameters as follows: degree cutoff = 2, node score cutoff = 0.6, k-core = 2, and max. depth = 100, and further cluster expansions were done by observing the network module visualization.

Enrichment analysis and functional interpretation of sub-modules and hub proteins were performed using the DAVID enrichment analysis tool (DAVID Bioinformatics Resources 6.8) and literature mining.

### 2.5. Identification and analysis of hub proteins

Intramodular hub proteins were selected according to the degree of each protein. The degree cutoff for hub selection was determined by analyzing the degree distribution and picking the top 10% of proteins with the highest degrees [20]. Furthermore, intermodular hubs were identified by analyzing the inter-modular connections. Among the proteins which connect different sub-modules, the proteins which connect at least 3 sub-modules were selected as intermodular hubs. Functional interpretations of hub proteins were performed by investigating the literature.

The methodology of this study is briefly illustrated in Fig.1.

**Fig.1.**
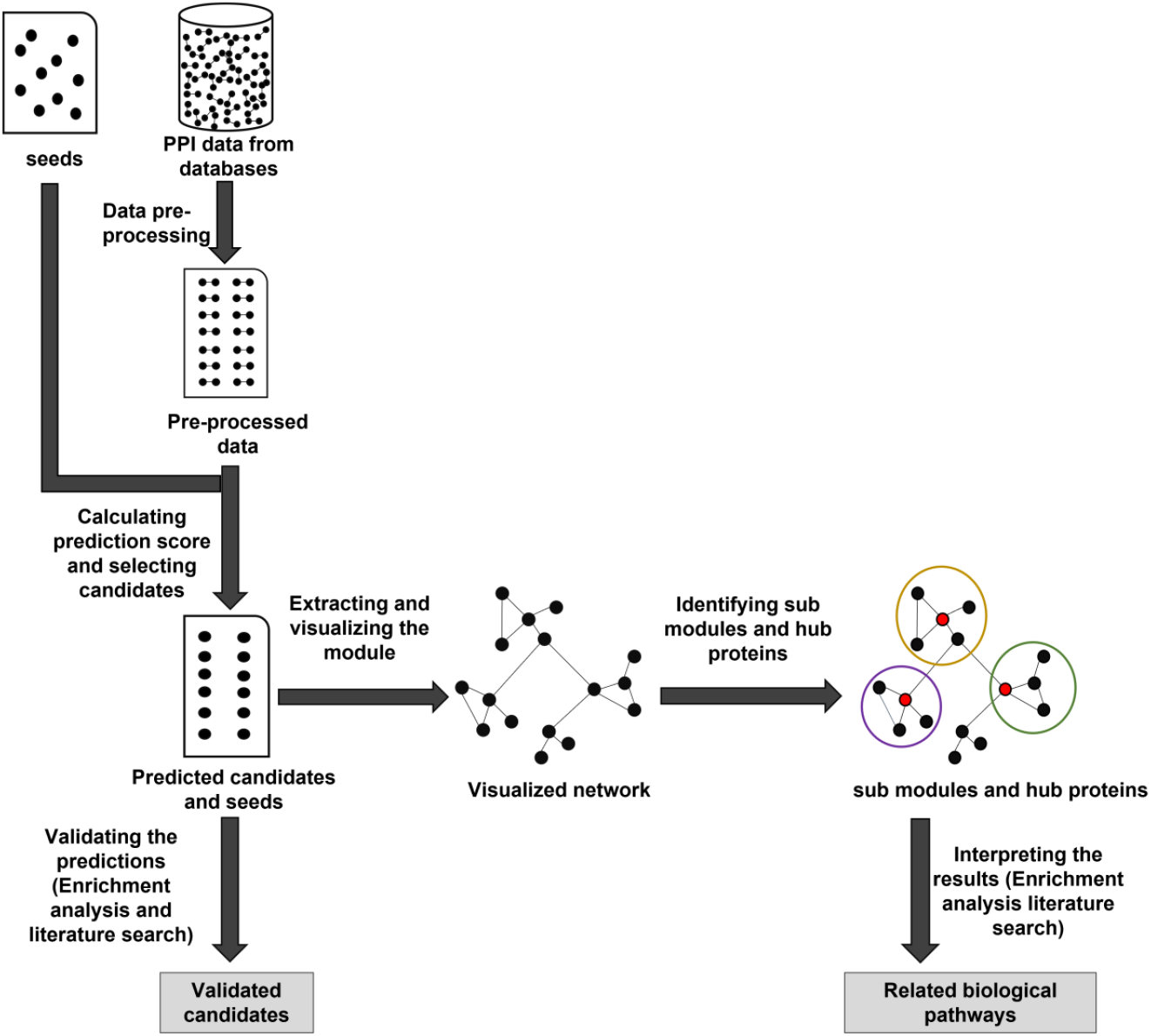
PPI network-based candidate protein prediction and validation workflow

All required scripts were written in python (version 2.7) and deposited in a GitHub repository: (https://github.com/Samadhi9/PPIN-analysis).

## 3. Results and Discussion

### 3.1. Data retrieval and data pre-processing

Altogether there were 51 seed proteins extracted from the literature and the STRING database (Supplementary Table A). The *O. sativa* STRING PPI network contained 25,106 proteins and 8,949,048 interactions. There were 21,212 proteins and 1,608,106 interactions after filtering by > 0.4 combined score cutoff. The number of interactions was reduced to 803,817 after removing duplicates.

### 3.2. Network-based candidate protein prediction and root development network module extraction

Hishigaki method was used for network-based candidate protein prediction, and after several trials and errors, the top 75 candidates were selected as the most suitable number of candidates for further analyses because it gave the best visualization of the root development network module by connecting most of the sub-modules. Moreover, a significant number of seed proteins were included in the extracted network module (Table 1).

**Table 1.** The number of seed proteins that were present and absent in the root development network modules according to different thresholds of selecting protein candidates.

|  | Top 20 candidates |  | Top 50 candidates |  | Top 75 candidates |  | Top 100 candidates |  |
| --- | --- | --- | --- | --- | --- | --- | --- | --- |
|  | Present | Absent | Present | Absent | Present | Absent | Present | Absent |
| No. of seeds | 38 | 13 | 41 | 10 | 45 | 6 | 45 | 6 |

As shown in Table 1, the top 20 and the top 50 candidates had a lower number of seeds present compared to the top 75 and 100. The top 75 and 100 had better seed retention and both retained the same number (45) of seed proteins.

Furthermore, degree distribution plots for the modules with 75 and 100 candidates showed an overlap with insignificant fluctuations (Fig. 2). Therefore, the top 75 proteins were selected focusing on the reliability of predictions and minimizing the data loss.

**Fig. 2.**
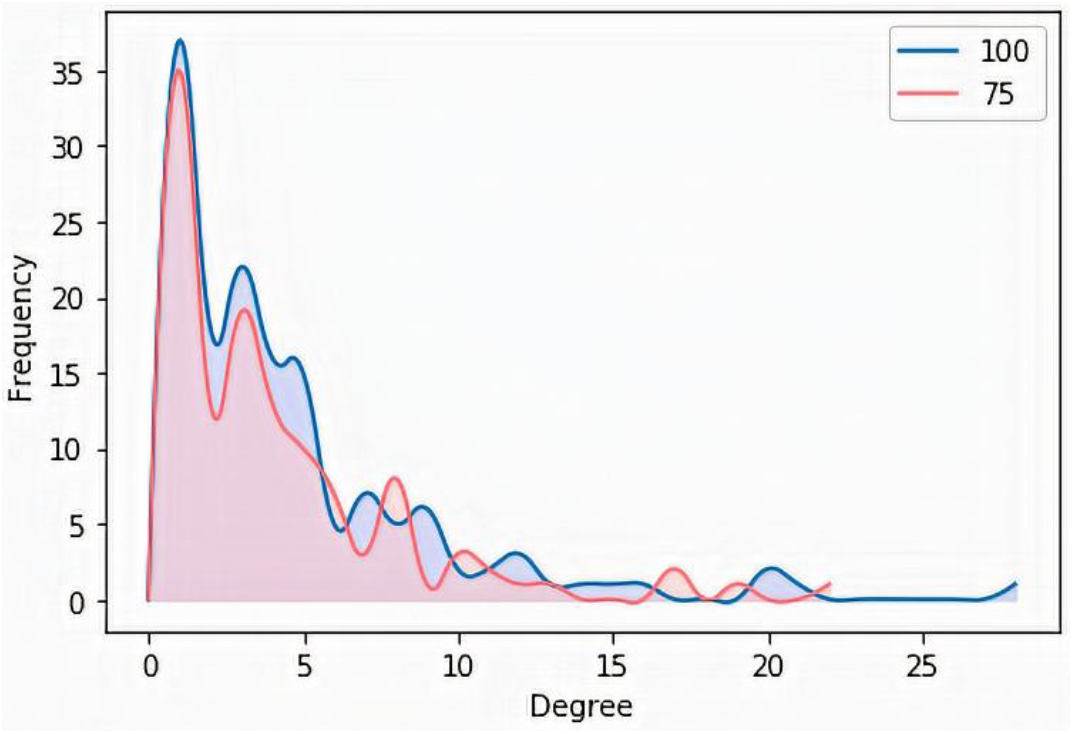
Degree distribution comparison between modules containing top 75 and 100 predicted proteins.

Although selecting the top 100 candidates connects more proteins in the network module, visualization was not clear due to congestion, and it did not reveal novel information about new sub-modules (Supplementary Fig. A). It was just an expansion of the existing sub-modules, which caused the integration of the 3^rd^ and the 4^th^ sub-modules shown in Fig.3. Although the sub-modules 3 and 4 are integrated as in Supplementary Fig. A, according to the enrichment analysis results, they are involved in different pathways: cytokinin-activated signaling pathway and cell wall organization, respectively. Therefore top 75 protein candidates were selected for further analyses. Their prediction scores are given in Supplementary Table 2.

**Fig. 3.**
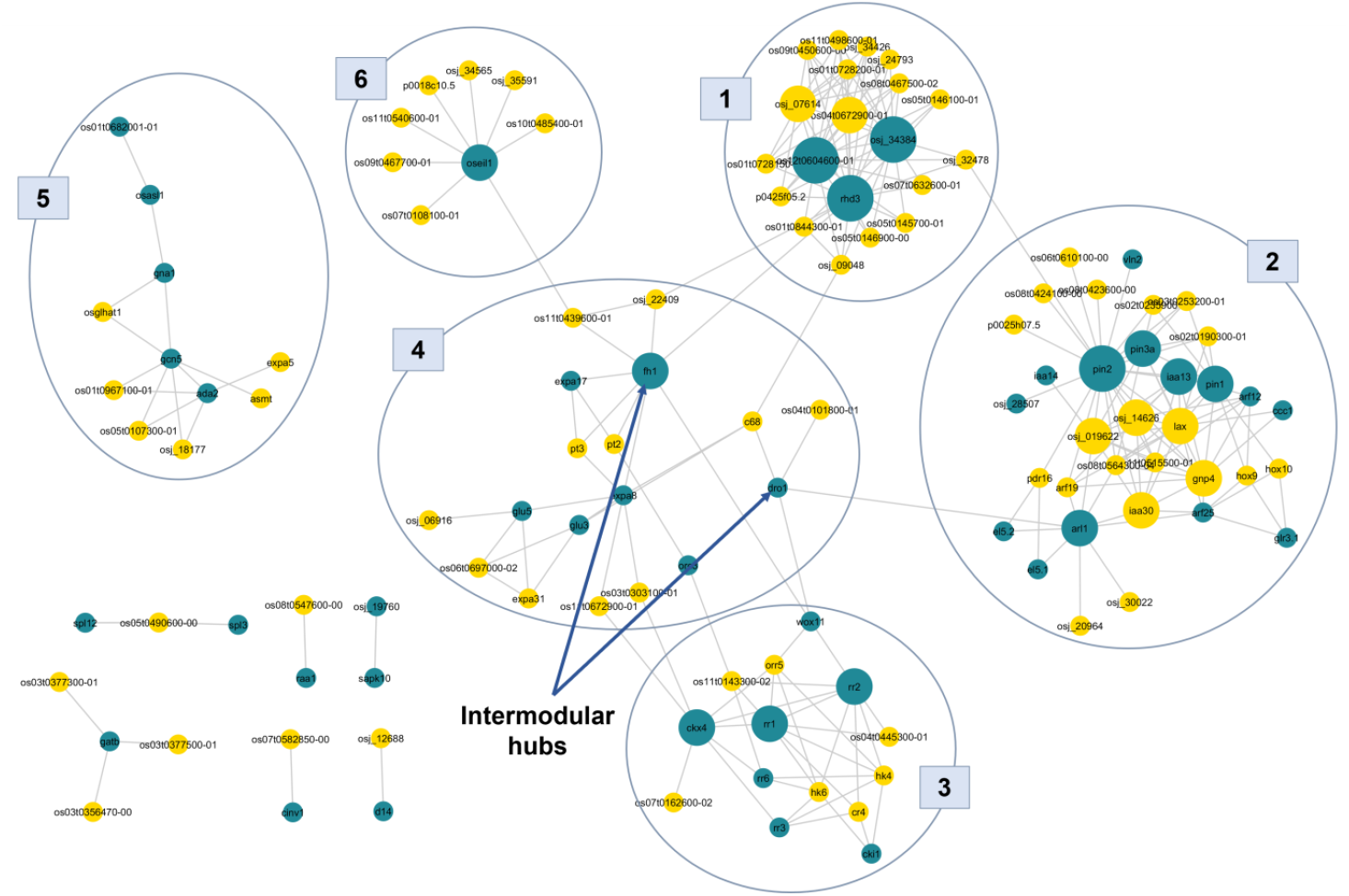
PPI network module for root development in *O. sativa* including the top 75 predicted proteins. Cyan nodes represent the seed proteins and yellow nodes represent the predicted proteins. Edges represent the interactions between proteins. The size of the node is proportional to the degree of the node. Large nodes represent a degree greater than or equal to 15 and medium nodes represent the hubs that have a degree within the 8-14 range. The numbered circles separate the sub-modules.

### 3.3. Root development network module visualization

Visualization of the root development PPI network module with the top 75 predicted proteins is given in Fig.3.

Altogether there were 120 proteins including 75 predicted candidates and 45 seed proteins (Supplementary Table B) (Fig. 3). However, 6 seed proteins (Supplementary Table C) were not visualized in the network because 3 of those were not included in the STRING raw dataset and 3 of those were removed while data pre-processing (Supplementary Table C).

### 3.4. Computational validation of the predictions

The enriched GO-BP terms from the enrichment analysis for the predicted root development protein candidates are given in Table 2

**Table 2.**
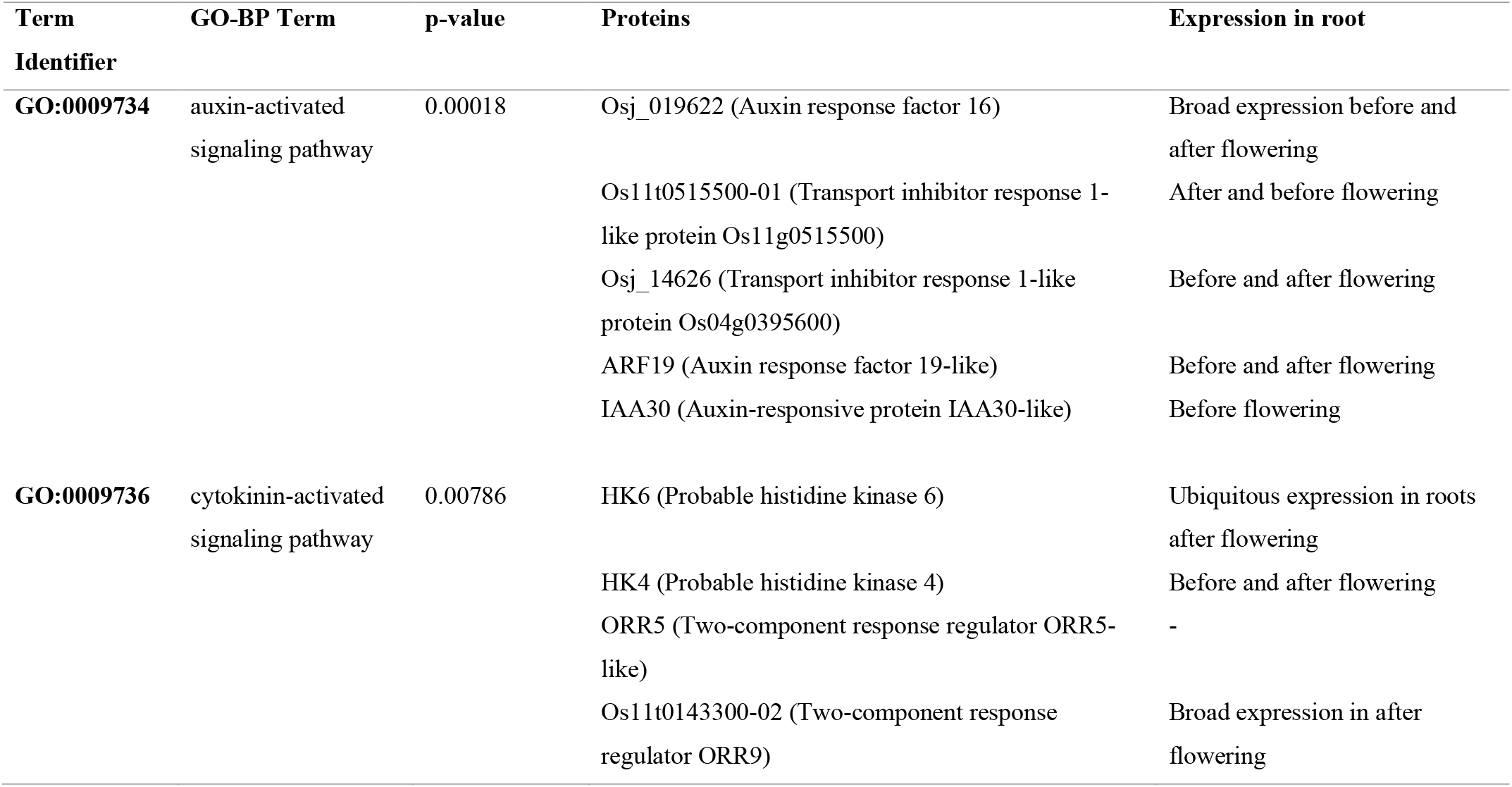
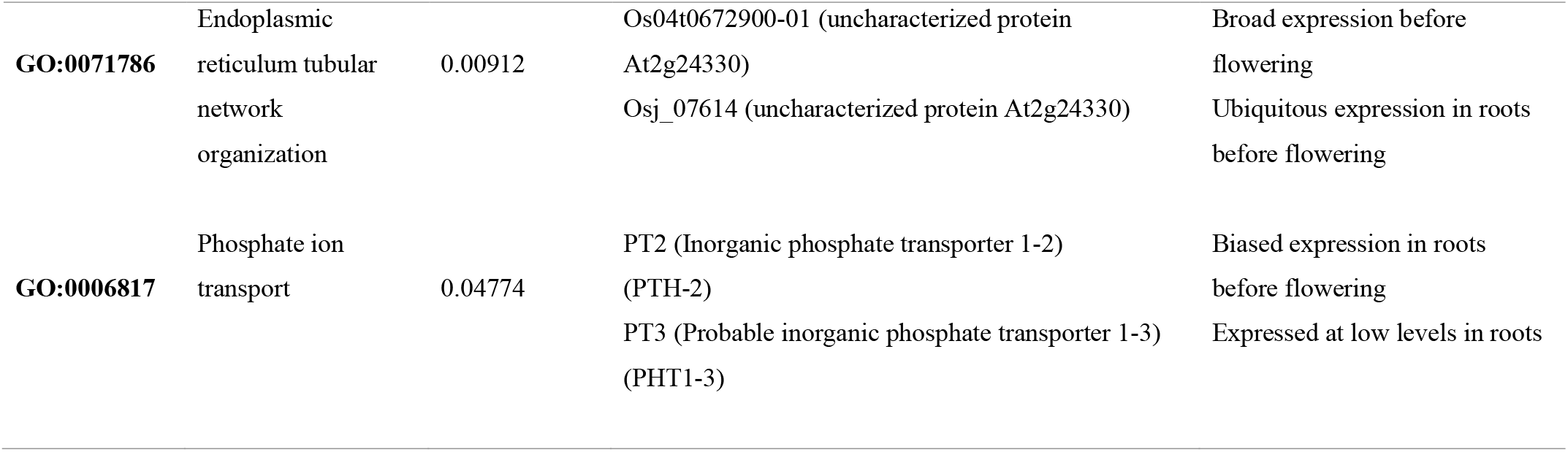
Enriched GO-BPs terms for predicted proteins which are related to root development and the pattern of expression of the corresponding genes in the root.

The GO term: auxin-activated signaling pathway (GO:0009734) is the most enriched term for the predicted protein list. Auxin is a growth coordinator hormone that regulates where, when, how much, and what sort of growth should occur in a plant. Auxins are expressed in many parts of the root such as root tip, root cap, root epidermis, etc. Moreover, auxins can be seen in the primary root, lateral roots, and root hairs. They are essential for many development processes in the root such as fine-tuning the growth rates, gravitropic root growth, cell division, proliferation, differentiation, and elongation of the root [38,39].

The biological process cytokinin-activated signaling pathway (GO:0009736) was also enriched according to Table 2. Cytokinin plays several roles in root development, including regulating the responses to the growth nutrients and the biotic and abiotic stresses. Furthermore, cytokinin regulates root differentiation, elongation, branching, and root architecture [39,40]. Moreover, it inhibits lateral root initiation and primary root elongation [40] while being essential for crown root development in *O. sativa* [4]. According to the enrichment analysis results, predicted proteins HK4, HK6, and ORR5 are in that pathway and their network neighbors (CR4, Os11t0143300-02, Os04t0445300-01) may also have a role in root development.

Results of enrichment analysis provide strong evidence to conclude that several predicted proteins are involved in the root development, and it is safe to speculate the predicted proteins as accurate predictions, which validates the prediction method.

The expression data in Table 2 were retrieved from the NCBI database, which was curated from the project: ‘Transcriptome profiling of various organs at different developmental stages in rice’ (BioProjectID: PRJNA243371). Root samples for the transcriptome profiling were taken before flowering and after flowering [37]

### 3.5. Identification and functional analysis of sub-modules

There were 6 identified sub-modules (Fig.3) in the root development PPI network module.

Enriched ontology terms for each sub-module were used to describe each sub-module. Proteins and enriched GO-BP terms for each sub-module are given below.

#### 3.5.1. Sub-module (1)

Tables 3 and 4 contain the proteins and the enriched GO-BPs for sub-module (1).

**Table 3.** NCBI gene descriptions, predicted status, and hub status of the 19 proteins in sub-module (1)

| **Name in network module** | **Gene description** | **predicted status** | **Hub/Non-hub** |
| --- | --- | --- | --- |
| **RHD3** | ROOT HAIR DEFECTIVE 3 | Seed | Hub |
| **Osj_34384** | ROOT HAIR DEFECTIVE 3 homolog 2 | Seed | Hub |
| **Os12t0604600-01** | ROOT HAIR DEFECTIVE 3 homolog 1 | Seed | Hub |
| **Os04t0672900-01** | uncharacterized protein At2g24330 | Predicted | Hub |
| **Osj_07614** | uncharacterized protein At2g24330 | Predicted | Hub |
| **Os08t0467500-02** | HVA22 like protein | Predicted | Non-hub |
| **Os09t0450600-00** | HVA22 like protein | Predicted | Non-hub |
| **Osj_24793** | HVA22-like protein k | Predicted | Non-hub |
| **Os01s0728150-00** | HVA22-like protein f | Predicted | Non-hub |
| **Os11t0498600-01** | HVA22-like protein e | Predicted | Non-hub |
| **P0425F05.2** | HVA22-like protein a | Predicted | Non-hub |
| **Os05t0146900-00** | uncharacterized | Predicted | Non-hub |
| **Osj_09048** | uncharacterized | Predicted | Non-hub |
| **Osj_32478** | uncharacterized | Predicted | Non-hub |
| **Os07t0632600-01** | uncharacterized | Predicted | Non-hub |
| **Os05t0145700-01** | uncharacterized | Predicted | Non-hub |
| **Os05t0146100-01** | uncharacterized | Predicted | Non-hub |
| **Os01t0844300-01** | peptidyl-prolyl cis-trans isomerase FKBP20-1 | Predicted | Non-hub |
| **Os01t0728200-01** | Discontinues | Predicted | Non-hub |

**Table 4.**
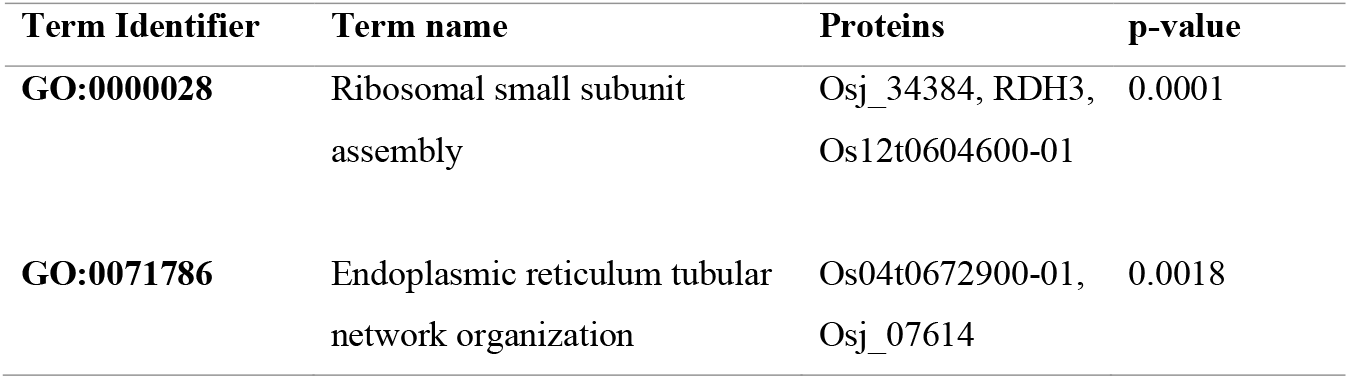
Significant enriched GO-BP terms from the enrichment analysis for sub-module (1)

| Term Identifier | Term name | Proteins | p-value |
| --- | --- | --- | --- |
| **GO:0000028** | Ribosomal small subunit assembly | Osj_34384, RDH3, Os12t0604600-01 | 0.0001 |
| **GO:0071786** | Endoplasmic reticulum tubular network organization | Os04t0672900-01, Osj_07614 | 0.0018 |

According to the results of the enrichment analysis in Table 4, it can be speculated that sub-module (1) is involved with the endoplasmic reticulum (ER) tubular network organization (GO:0071786). The ER tubular network is involved with root hair tip growth [41]. Though the hub protein RHD3 was not listed under the ER tubular network organization (Table 4), it involves root hair tip growth by organizing the ER tubular network. [41,42]. Also, *Arabidopsis rhd3* mutant causes short and wavy root hairs [43]. Furthermore, HVA22d protein co-localizes with RHD3 and involves ER shaping [42], and several proteins of the HVA22 family (protein HVA22 like protein, HVA22-like protein k, HVA22-like protein f, HVA22-like protein e, and HVA22-like protein a), which were predicted, can be seen under this sub-module. According to this information, predicted HVA22 family proteins can be speculated to be involved with root development in *O. sativa*.

Proteins Osj_34384, Os12t0604600-01, Os04t0672900-01, and Osj_07614 were also recognized as hubs in this sub-module. Among these, proteins Os04t0672900-01 and Osj_07614 are two predicted candidates which have not been characterized yet. According to the results, they probably have important roles in root development.

#### 3.5.2. Sub-module (2)

Tables 5 and 6 contain the proteins and the enriched GO-BPs for sub-module (2), respectively.

**Table 5.** NCBI gene descriptions, predicted status, and hub status of the 34 proteins in sub-module (2)

| **Name in network module** | **Gene description** | **predicted status** | **Hub/Non-hub** |
| --- | --- | --- | --- |
| **ARL1** | LOB domain-containing protein 29 | Seed | Hub |
| **PIN1** | auxin efflux carrier component 1 | Seed | Hub |
| **PIN2** | probable auxin efflux carrier component 2 | Seed | Hub |
| **PIN3A** | probable auxin efflux carrier component 3a | Seed | Hub |
| **IAA13** | auxin-responsive protein IAA13 | Seed | Hub |
| **EL5.2** | E3 ubiquitin-protein ligase EL5 | Seed | Non-hub |
| **EL5.1** | E3 ubiquitin-protein ligase EL5 | Seed | Non-hub |
| **ARF25** | auxin response factor 25 | Seed | Non-hub |
| **ARF12** | auxin response factor 12 | Seed | Non-hub |
| **Osj_28507** | auxin transport protein BIG | Seed | Non-hub |
| **CCC1** | cation-chloride cotransporter 1 | Seed | Non-hub |
| **IAA14** | auxin-responsive protein IAA14 | Seed | Non-hub |
| **GLR3.1** | glutamate receptor 3.1 | Seed | Non-hub |
| **VLN2** | new submission | Seed | Non-hub |
| **Osj_019622** | auxin response factor 16 | Predicted | Hub |
| **IAA30** | auxin-responsive protein IAA30 | Predicted | Hub |
| **GNP4** | protein LAX PANICLE 2 | Predicted | Hub |
| **LAX1** | transcription factor LAX PANICLE 1 | Predicted | Hub |
| **Osj_14626** | transport inhibitor response 1-like protein<br>Os04g0395600 | Predicted | Hub |
| **HOX10** | homeobox-leucine zipper protein HOX10-like | Predicted | Non-hub |
| **HOX9** | homeobox-leucine zipper protein HOX9 | Predicted | Non-hub |
| **ARF19** | auxin response factor 19 | Predicted | Non-hub |
| **Os02t0190300-01** | putative multidrug resistance protein | Predicted | Non-hub |
| **Os02t0235900-01** | tetrahydrocannabinolic acid synthase | Predicted | Non-hub |
| **Os03t0253200-01** | serine/threonine-protein kinase WAG1 | Predicted | Non-hub |
| **Os06t0610100-00** | Discontinued | Predicted | Non-hub |
| **Os08t0423600-00** | alpha carbonic anhydrase 7-like | Predicted | Non-hub |
| **Os08t0424100-00** | alpha carbonic anhydrase 7 | Predicted | Non-hub |
| **Os08t0564300-04** | ABC transporter B family member 1 | Predicted | Non-hub |
| **Os11t0515500-01** | transport inhibitor response 1-like protein<br>Os11g0515500 | Predicted | Non-hub |
| **Osj_20964** | uncharacterized | Predicted | Non-hub |
| **Osj_30022** | disease resistance protein RPM1 | Predicted | Non-hub |
| **P0025H07.5** | alpha carbonic anhydrase 7 | Predicted | Non-hub |
| **PDR16** | ABC transporter G family member 32-like | Predicted | Non-hub |

**Table 6.**
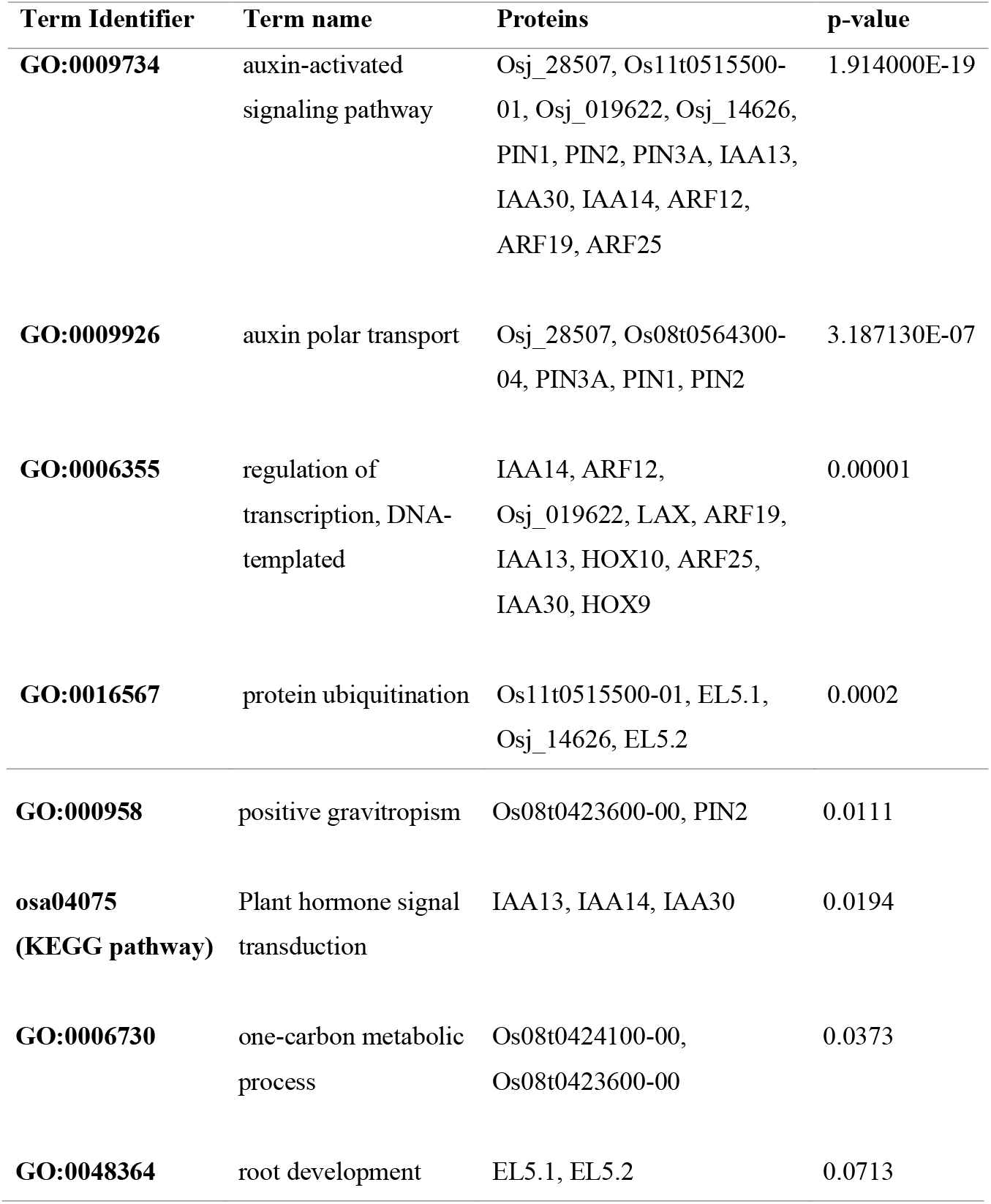
Selected enriched GO-BPs and KEGG pathways from the enrichment analysis for sub-module (2)

| **Term Identifier** | **Term name** | **Proteins** | **p-value** |
| --- | --- | --- | --- |
| **GO:0009734** | auxin-activated signaling pathway | Osj_28507, Os11t0515500-01, Osj_019622, Osj_14626, PIN1, PIN2, PIN3A, IAA13, IAA30, IAA14, ARF12, ARF19, ARF25 | 1.914000E-19 |
| **GO:0009926** | auxin polar transport | Osj_28507, Os08t0564300-04, PIN3A, PIN1, PIN2 | 3.187130E-07 |
| **GO:0006355** | regulation of transcription, DNA-templated | IAA14, ARF12, Osj_019622, LAX, ARF19, IAA13, HOX10, ARF25, IAA30, HOX9 | 0.00001 |
| **GO:0016567** | protein ubiquitination | Os11t0515500-01, EL5.1, Osj_14626, EL5.2 | 0.0002 |
| **GO:000958** | positive gravitropism | Os08t0423600-00, PIN2 | 0.0111 |
| **osa04075**<br>(KEGG pathway) | Plant hormone signal<br>transduction | IAA13, IAA14, IAA30 | 0.0194 |
| **GO:0006730** | one-carbon metabolic<br>process | Os08t0424100-00,<br>Os08t0423600-00 | 0.0373 |
| **GO:0048364** | root development | EL5.1, EL5.2 | 0.0713 |

According to Table 6, the most enriched function for the sub-module (2) is the auxin-activated signaling pathway (GO:0009734), and most of the other functions of sub-module (2) are also related to the phytohormone auxin. The auxin signaling pathway combines transport inhibitor response 1 (TIR1), auxin response factors (ARFs), and auxin/indole acetic acid (AUX/IAA) transcriptional repressors together [39]. According to the enrichment analysis results, hub proteins Osj_019622, Osj_14626, PIN1, PIN2, PIN3A, IAA13, and IAA30, and non-hub proteins Osj_28507, Os11t0515500-01, IAA14, ARF12, ARF19, and ARF25 in sub-module (2) participate in the auxin signaling pathway.

Auxin polar transport (GO:0009926) is another enriched function for the sub-module (2) which includes PIN2, PIN1, PIN3A, Osj_28507, and Os08t0564300-04 as annotated proteins. The protein PIN2, a hub with the highest degree of 22, is a potential candidate gene for improving root system architecture in *O. sativa* [44,45], and PIN1 and PIN3A are also hubs, which are central to the stability of this sub-module.

Auxin polar transport is regulated by the PIN-FORMED (PIN) efflux carriers. PIN polarity plays a crucial role in developing proper organs and proliferation in root proximal meristem [38,39,46]. For example, the intramodular hub protein PIN2 works for root development by positioning and emerging root hairs [38], and according to Inahashi, et al., [47], the *OsPIN2* gene regulates the seminal root elongation and lateral root formation in *O. sativa*. Moreover, overexpression of the *OsPIN2* significantly decreases the number of adventitious roots and the total root length by 22%–28% [44]. Furthermore, Overexpression of the gene *OsPIN3a* has led to the development of longer roots and more adventitious roots [48], and hang et al., [48] suggest that crown root development is controlled by auxin signaling through PIN proteins. Another member of this sub-module, the protein PIN1, is also a hub protein, and overexpression of gene *OsPIN1* increases the emergence of adventitious roots, the primary root length, and the number of lateral roots [46,49].

The roots have the ability to change their growing orientation in response to the changes in gravity [38,45], and it is controlled by the asymmetric distribution of auxin at the root tip. PIN family, AUX1 (AUXIN-INSENSITIVE1), and other members of the auxin transport pathway contribute to this auxin distribution [38,45]. Deletion in the *OsPIN2* gene has displayed gravitropic root growth phenotypes, as shootward auxin distribution in the lower side of the root is largely repressed during a gravity stimulus by the mutation of *OsPIN2* [44,45]. This shows that PIN2 is also essential for root gravitropism. Moreover, the crown root growth angle is an important component of the *O. sativa* roots, and the *OsPIN2* mutant, *lra1* has displayed larger root angles [45]. This indicates that the protein PIN2 is important in regulating crown root growth angle. Furthermore, *OsPIN3a has* shown a notable up-regulation in *OsPIN2* mutant *lra1* since *OsPIN3a* compensates for the loss of *OsPIN2* (agravitropic root phenotype) to some extent [45]. One of the enriched terms for the sub-module (2) is positive gravitropism (GO:0009958), and the seed protein PIN2 and the predicted candidate Os08t0423600-00 have been annotated to that process. This evidence proves that the majority of the sub-module 2 proteins contribute to root development via auxin regulation.

#### 3.5.3. Sub-module (3)

Tables 7 and 8 contain the proteins and the enriched GO-BPs for sub-module (3), respectively.

**Table 7.** NCBI gene descriptions, predicted status, and hub status for the 14 proteins in sub-module (3)

| **Name in network module** | **Gene description** | **predicted status** | **Hub/ Non-hub** |
| --- | --- | --- | --- |
| **RR1** | two-component response regulator ORR1 | Seed | Hub |
| **RR2** | two-component response regulator ORR2 | Seed | Hub |
| **CKX4** | cytokinin dehydrogenase 4 | Seed | Hub |
| **RR3** | two-component response regulator ORR3 | Seed | Non-hub |
| **RR6** | two-component response regulator ORR6 | Seed | Non-hub |
| **CKI1** | Casein kinase 1 | seed | Non-hub |
| **WOX11** | WUSCHEL-related homeobox 11 | Seed | Non-hub |
| **CR4** | putative receptor protein kinase CRINKLY4 | Predicted | Non-hub |
| **HK4** | probable histidine kinase 4 | Predicted | Non-hub |
| **HK6** | probable histidine kinase 6 | Predicted | Non-hub |
| **ORR5** | two-component response regulator ORR6 | Predicted | Non-hub |
| **Os11t0143300-02** | two-component response regulator ORR9 | Predicted | Non-hub |
| **Os04t0445300-01** | uncharacterized | Predicted | Non-hub |
| **Os07t0162600-02** | probable carboxylesterase | Predicted | Non-hub |

**Table 8.** Selected enriched GO-BPs and KEGG pathways of enrichment analysis for sub-module (3)

| **Term Identifier** | **Term name** | **Protein** | **p-value** |
| --- | --- | --- | --- |
| **GO:0009736** | cytokinin-activated signaling pathway | HK4, HK6, RR2, RR6, RR1, RR3, ORR5 | 1.166273E-12 |
| **GO:0000160** | phosphorylation signal transduction system | RR2, RR6, RR1, RR3, ORR5 | 5.924492E-08 |
| **osa04075(KEGG pathway)** | Plant hormone signal transduction | RR2, RR6, RR1, RR3, ORR5 | 0.00002 |
| **GO:0006355** | regulation of transcription, DNA-templated | RR2, RR6, RR1, RR3, ORR5, WOX11 | 0.00045 |
| **GO:0006468** | protein phosphorylation | HK4, HK6, CR4 | 0.0163 |

According to Table 8, the top enriched GO-BP term for sub-module (3): Cytokinin-activated signaling pathway (GO:0009736), is mediated by a two-component system, and the signaling is transmitted by transcription activators and repressors in a phosphorylation signal transduction system (GO:0000160) [50,51]. The two-component system comprises three functional modules: sensory histidine kinase (HK), histidine phosphor transfer protein (HP), and response regulator (RR). Cytokinins are sensed by membrane-located HK receptors that transmit signals via HPs to nuclear RRs that activate or repress transcription [52].

The phytohormone cytokinin is present in the *O. sativa* root [43] and participates in regulating root development and root architecture as described in section 3.4. This indicates the involvement of the Cytokinin-activated signaling pathway in *O. sativa* root development and involvement of sub-module (3) proteins in root development through the Cytokinin-activated signaling transduction pathway.

#### 3.5.4. Sub-module (4)

Tables 9 and 10 contain the proteins and the enriched GO-BPs in sub-module (4).

**Table 9.** NCBI gene descriptions, predicted status, and hub status for the 18 Proteins in sub-module (4)

| **Name in network module** | **Gene description** | **predicted status** | **Hub/ Non-hub** |
| --- | --- | --- | --- |
| **FH1** | formin-like protein 1 | Seed | Hub |
| **DRO1** | uncharacterized | Seed | Hub |
| **GLU3** | endoglucanase 12 | Seed | Non-hub |
| **GLU5** | endoglucanase 2 | Seed | Non-hub |
| **ORC3** | origin of replication complex subunit 3 | Seed | Non-hub |
| **EXPA8** | expansin-A8 | Seed | Non-hub |
| **EXPA17** | expansin-A17-like | Seed | Non-hub |
| **C68** | probable LRR receptor-like serine/threonine-protein kinase At5g45780 | Predicted | Non-hub |
| **EXPA31** | expansin-A31 | Predicted | Non-hub |
| **Os03t0303100-01** | serine/arginine repetitive matrix protein 2 | Predicted | Non-hub |
| **Os04t0101800-01** | uncharacterized | predicted | Non-hub |
| **Os06t0697000-02** | probable xyloglucan endotransglucosylase/hydrolase protein 25 | Predicted | Non-hub |
| **Os11t0439600-01** | probable apyrase 3 | Predicted | Non-hub |
| **Os11t0672900-01** | serine/arginine repetitive matrix protein 1 | Predicted | Non-hub |
| **Osj_06916** | uncharacterized | Predicted | Non-hub |
| **Osj_22409** | COBRA-like protein 10 | Predicted | Non-hub |
| **PT2** | inorganic phosphate transporter 1-2 | Predicted | Non-hub |
| **PT3** | probable inorganic phosphate transporter1-3 | predicted | Non-hub |

**Table 10.** Selected enriched GO-BPs of enrichment analysis for sub-module (4)

| **Term Identifier** | **Term name** | **Proteins** | **p-value** |
| --- | --- | --- | --- |
| **GO:0071555** | cell wall organization | GLU5, GLU3, Os06t0697000-02 | 0.0091 |
| **GO:0006817** | phosphate ion transport | PT2, PT3 | 0.0129 |
| **GO:0030245** | cellulose catabolic process | GLU5, GLU3 | 0.0185 |

As given in Table 10, sub-module (4) has proteins that are associated with both plant cell wall organization (GO:0071555) and cellulose catabolic process (GO:0030245) pathways. The proteins GLU5, GLU3, and Os06t0697000-02 involve cell wall organization, and GLU5 and GLU3 involve the cellulose catabolic process. Cell walls are important in any form of plant development, and cellulose is the major component of the plant cell wall. Cellulose biosynthesis and cell wall loosening enable turgor-driven cell expansion in growing plants, and it has been speculated that endo-1,4-b-glucanases (EGases) play a central role in these complex activities [53]. GLU3 and GLU5 (GLUs) have been directly annotated to the hydrolysis of endo-1,4-b-glucanases [54,55]. GLUs play important roles in root development and *glu* mutants have reduced root development [55]. Furthermore, GLU5 is expressed in lateral root primordia during auxin-induced lateral root development [54].

Phosphate ion transport (GO:0006817) in a plant is mediated by several transporter protein families such as the Pht1 family [56]. PT2 and PT3 belong to the pht1 family and are predicted candidates in sub-module (4). Both root hair length and frequency increase in response to phosphate (Pi) starvation and the gene expression of *OsPT2* is increased during Pi starvation. Therefore, it is reasonable to speculate that PT2, which is a predicted protein, probably has a direct association with Pi transport. [56,57].

FH1 is an intramodular hub in sub-module (4) and also an intermodular hub. FH1 has been directly annotated to root hair development [58,59]. EXPA8 in sub-module 4 (degree = 7) was not considered as a hub according to the hub selection criteria of this study. However, it is a root-specific expansin protein, and expansins are plant cell wall proteins that are involved in cell wall modifications [60,61]. Overexpression of the *OsEXPA8* gene has shown improved root system architecture with longer primary roots and a higher number of lateral roots and root hairs [60,61]. Moreover, repression of *OsEXPA8* has reduced the cell size of the root vascular system and plant height [61]. This evidence prove that Sub-module (4) is linked to the root development via cell wall organization.

Proteins of other sub-modules are listed in Supplementary Table D. They did not have any significant enriched GO-BP terms related to root development and need further investigation to confirm their involvement in root development.

### 3.6. Identification and analysis of Hub proteins

There are two types of hubs: intramodular and intermodular hubs [21]. Intramodular hubs are found within a functional module while intermodular hubs act between the modules to interconnect them [20–22].

#### 3.6.1. Intramodular hubs

For this study, the top 10% of proteins with the highest degrees were selected [62], which corresponds to a degree cutoff of 8, resulting in 20 proteins as intramodular hubs (Table 11).

**Table 11.** Details of intramodular hub proteins

| **Protein** | **Degree** | **Type**<br>**(seed/predicted)** |
| --- | --- | --- |
| **PIN2** | 22 | Seed |
| **RHD3** | 19 | Seed |
| **Osj_34384** | 17 | Seed |
| **Os12T0604600-01** | 17 | Seed |
| **LAX1** | 13 | Predicted |
| **GNP4** | 12 | Predicted |
| **Os04T0672900-01** | 11 | Predicted |
| **Osj_07614** | 11 | Predicted |
| **Osj_019622** | 10 | Predicted |
| **ARL1** | 10 | Seed |
| **PIN1** | 10 | Seed |
| **PIN3A** | 9 | Seed |
| **IAA30** | 8 | Predicted |
| **Osj_14626** | 8 | Predicted |
| **FH1** | 8 | Seed |
| **IAA13** | 8 | Seed |
| **CKX4** | 8 | Seed |
| **RR2** | 8 | Seed |
| **RR1** | 8 | Seed |
| **OSEIL1** | 8 | Seed |

Some of the hubs are from the seed proteins and others are predicted candidates for root development in *O. sativa*. The predicted hubs are important findings because their relevance in root development has not been revealed to date. They also confirm the accuracy and the importance of the predictions. Among the predicted hubs, there were candidates annotated to the enriched GO-BP terms related to the root development in *O. sativa*, although they lack direct experimental evidence for root development. For example, *Os04t0672900-01 and Osj_07614* of sub-module (1) annotated to GO:0071786: Endoplasmic reticulum tubular network organization from Table 4, which is a GO-BP term associated with root development.

Also, transcription factors LAX1 (LAX PANICLE 1) and GNP4 (LAX PANICLE 2) are predicted candidates, which were identified as hubs. In *O. sativa*, LAX1 and GNP4 are required for the formation of Axillary meristem throughout the plant’s lifespan [63,64]. Also, LAX1 shows non-cell-autonomous action (mutant extends beyond the mutant cells); however, its molecular basis has not been revealed yet [63]. Although the functions of lax genes in *O. sativa* panicle have been studied, their functions in the root are yet to be revealed. The above results provide evidence for their involvement in root development. Therefore, this study provides potential candidates for selecting important proteins for future *O. sativa* root development studies.

The seed protein OSEIL1 is an intramodular hub and is involved in root development in *O. sativa*. It is a transcription factor participating in the ethylene signaling pathway, which promotes *O. sativa* root elongation [65]. Most importantly, OSEIL1 connects with 8 predicted candidates and joins the sub-module (4) and (6) together (Fig.3). Therefore, according to our results, OSEIL1 is a critical protein for root development.

#### 3.6.2. Intermodular hubs

Intermodular hubs connect different sub-modules (Fig.3) and are important in linking the different metabolic/biological pathways. Two intermodular hubs were detected in this study (Table 12).

**Table 12.** Details of intermodular hub proteins

| **Intermodular hub** | **Degree** | **Type**<br>**(seed/predicted)** | **Connected proteins** | **Sub-module**<br>**(Fig. 3)** |
| --- | --- | --- | --- | --- |
| **DRO1**<br>**(In 4<sup>th</sup> sub-module)** | 4 | Seed | C68,<br>Os04t0101800-01 | 4 |
|  |  |  | ARL1 | 2 |
|  |  |  | WOX11 | 3 |
| **FH1**<br>**(In 4<sup>th</sup> sub-module)** | 8 | Seed | PT3, PT2, EXPA8,<br>EXPA17,<br>Osj_22409,<br>Os11t0439600-01 | 4 |
|  |  |  | RHD3 | 1 |
|  |  |  | WOX11 | 3 |

As shown in Table 12, the protein DRO1 (DEEPER ROOTING 1) in sub-module 4 works as an intermodular hub and connects sub-modules 2, 3, and 4. Therefore, disturbance to the DRO1 can potentially disrupt the interconnectivity of the pathways or the proper mechanism of those sub-modules. Analysis of DRO1 using iDNA-Prot (http://www.jci-bioinfo.cn/iDNA-Prot) [66], which is a web tool for identifying DNA binding domains in proteins [66], revealed that it may be a DNA binding protein. Highly expressed *DRO1* gene is involved in the regulation of deep rooting by increasing root growth angle which promotes the root growth in a more downward direction [67]. Furthermore, *DRO1* enhances nitrogen uptake and cytokinin fluxes at the late stages of development by deep rooting which resulted in a high yield in *O. sativa*. Therefore, *DRO1* can be used to develop *O. sativa* cultivars that have high yields under both drought and non-drought conditions by controlling the root system architecture [67,68]. As shown in this study, the intermodular hub DRO1 plays a major role in interconnecting and potentially regulating the 3 submodules of root development and can be a valuable candidate for further experimental studies.

The FH1 (formin-like protein 1), which is in the sub-module (4), is an intramodular hub and a critical regulator of the *O. sativa* root hair development. *Osfh1* mutant exhibited growth defects on root hairs. These defects depend on the environmental conditions and were only exhibited when roots were submerged in a solution [58]. According to Huang et al., [59], the external supplies of the nutrients or the hormones could not rescue the defective mutant. Therefore, FH1 is a crucial protein for the growth of *O. sativa* as rice is grown under water-logged conditions in the field until the fruit ripening stage. [58,59].

FH1 is also identified as an intermodular hub and it connects sub-modules (1), (3), and (4) (Table 12, Fig 3). Formins regulate the growth and elongation of the root hairs and cell wall loosening and synthesis, which are required for root hair development [58]. Furthermore, the sub-module (4) which is annotated to the cell wall organization, and the sub-module (1) which is mainly recognized for root hair development are connected by FH1. This functional significance of FH1 is also confirmed by our results, which represent it as an intermodular hub. Since it is connected with 3 sub-modules, it could have more roles in different pathways which are not yet revealed.

### 3.7. Conclusion

In this study, we analyzed the network structure of root development proteins, during which 75 new protein candidates, 6 sub-modules, 20 intramodular hubs, and 2 intermodular hubs were identified. Validation of predictions and analysis of hubs and sub-modules justified that some predictions are annotated to the biological processes associated with root development, which confirmed the accuracy of the predictions. Finding hub proteins was one of the important outcomes of this study, which enabled identifying the important module members central to the stability of the root development module in *O. sativa*. All the above information was gathered in a computational setting in a single analysis, which consumed far less time and resources than wet-lab experiments (because 75 important protein candidates were extracted from 25,106 proteins). This opened new directions for future wet lab and dry lab studies. To our knowledge, this is the first study that analyzes the PPI network module for root development in *O. sativa*. Therefore, these findings are the first to show the PPI interaction structure underlying root development, which depicts the importance and applicability of network analysis on other plant phenotypes as well.

### 3.8. Future studies

This study led to the prediction of proteins important for root development, such as Os04t0672900-01, Osj_07614, and DRO1, which are not yet characterized. *In-silico* methods can be used for the characterization through the study of homolog and ortholog domain analysis and 3D structure prediction. Further analysis of identified hub proteins, especially their functions and the involvement in the metabolic pathways, will clarify the molecular mechanism of root development. Moreover, wet lab experiments can be conducted to validate these predictions and confirm the biological role of each protein in root development. Important hub proteins identified through these methods can be used for the genetic improvement of *O. sativa*, such as for drought resistance and submergence tolerance, through genetic breeding and engineering.

## List of Abbreviations

PPI: Protein-Protein Interactions
IAA: Indole-3-Acetic Acid
NAA: Naphthaleneacetic Acid
IDs: Identifiers
GO: Gene Ontology
GO-BP: Biological Process component of the Gene Ontology
ER: Endoplasmic Reticulum
PIN: PIN-FORMED
HK: Histidine Kinase
HP: Histidine Phosphor transfer
RR: Response Regulator
Pi: Phosphate
FH2: Formin Homology-2
PR: Primary Root
CR: Crown Root
ARF: Auxin Response Factor
EGases: Endo-1,4-b-glucanase

## Authors’ contributions

All authors planned and designed the experiments. SW wrote the Python scripts for the analysis and performed the experiments under the supervision of ST and PF. All authors analyzed the results, read, and approved the final manuscript

