## Supplemental Files for "Protein-protein interaction (PPI) network analysis reveals important hub proteins and sub-network modules for root development in rice (*Oryza sativa*)"

### Appendices

*Tables*

**Supplementary Table A**: Details of seed proteins

| Protein | Specific location | Involvement in root development | Function | Source |
|---|---|---|---|---|
| **ada2** | Primary root (PR), crown root (CR) | Root elongation of PR and CR, increase number of CR | Auxin or cytokinin signaling | Literature |
| **aim1** | Root meristem | Root growth | Salicylic acid biosynthesis promotes reactive oxygen species accumulation | Literature |
| **arf1** | Adventitious root | Root formation | Auxin response factor, transcriptional factor | Literature |
| **arf12** | Primary root | Longer primary roots | Auxin response factor, transcriptional factor | Literature |
| **arf25** | Crown root | Root development | Auxin response factor, transcriptional factor | Literature |
| **arl1** | Crown root, lateral root, adventitious root | Root formation, root development | Auxin-mediated cell de-differentiation | Literature |
| **crl4/gnom1** | Crown root | Root initiation | Polar auxin transportation | Literature |
| **ccc1** | Root tip | Root elongation | Osmotic regulation | Literature |
| **cinv1** | Primary root, lateral root | Root elongation, root formation | Cleaves sucrose into glucose and fructose, floral transition, and pollen development | STRING DB |
| **cki1** | Lateral root, adventitious root, primary root | Root elongation, increase the number of roots. | Involved in ABA and brassinosteroid signaling pathways. | STRING DB |

| | | | | |
|---|---|---|---|---|
| **ckx4** | Crown root | Root formation, root emergence, and development increase the number of crown roots. | CK signaling pathway | Literature |
| **cinv1 (cyt-inv1)** | - | Root elongation | Mutant accumulates sucrose and had reduced levels of hexose | Literature |
| **d14** | Crown roots | Root elongation | Involved in the strigolactone signaling pathway | Literature |
| **DRO1** | Crown roots | Increases root angle and facilitate root growth more downward | enhances nitrogen uptake and cytokinin fluxes | Literature |
| **el5.1/el5.2** | Root primordia | Maintains cell viability after the initiation of root primordial formation, | mediate the degradation of cytotoxic proteins produced in root cells | STRING DB |
| **expa17** | Root hair | Root elongation | Cell wall remodeling | Literature |
| **expa8** | Root and shoot | Root elongation, fewer LRs, and short root hairs | Encodes for cell wall localized protein | Literature |
| **fh1** | Root hair | Root elongation | Actin-binding protein | Literature |
| **gatb** | Primary root, root tip | Growth, cell division, and elongation | Maintaining mitochondrial structure and function | Literature |
| **gcn5** | Primary root, crown root | Root elongation of PR and CR, increase the number of CR | Auxin or cytokinin signaling | Literature |
| **glr3.1** | Root meristem (apical) | Root elongation, maintenance of cell division, survival in | Glutamate-gate receptor | Literature |

| | | | | |
|---|---|---|---|---|
| | | the root meristem in early seedlings | | |
| **glu3** | - | Root elongation | Encodes for putative membrane-bound endo-1,4-β-glucans. Mutant lower the cellulose contents in root cell walls | Literature |
| **glu5** | Lateral root development | Root development | Endohydrolysis of (1->4)-beta-D-glucosidic linkages in cellulose | STRING DB |
| **gna1** | Root elongation zone | Root elongation, maintaining normal root cell shape | Plays an important role in protein and lipid glycosylation | Literature |
| **iaa13** | Lateral root | Root formation | Member of IAA gene family, transcriptional factor | Literature |
| **iaa14** | Lateral root | Root initiation | Member of IAA gene family, transcriptional factor | Literature |
| **mst3** | Root system | Root development | Sugar transporter | STRING DB |
| **mt2c** | Lateral root | Root development, root initiation | Seed embryo germination by regulating CK level, reactive oxygen species scavenger in the cytosol | STRING DB |
| **orc3** | Lateral root | Root development | Component of the origin recognition complex that binds origins of replication | Literature |
| **os01t0682001-01** | Seedling roots | Primary ammonium ions assimilation | Reutilization of glutamine in | STRING DB |

| | | | | |
|---|---|---|---|---|
| | | | developing organs. Plays a role in the development of tillers | |
| **os12t0604600-01** | Root hair | Root development | Probable GTP-binding protein | STRING DB |
| **osasl1** | Primary root, root | Primary root elongation, normal root growth | Arginine biosynthesis | Literature |
| **oseil1** | - | Root elongation | Involved in the ethylene signaling pathway, transcriptional factor | Literature |
| **osj_19760** | Root hair | Form shorter root hairs | Reduced ABA signaling | Literature |
| **osj_28507** | Lateral root | Influencing auxin-mediated developmental responses root production (ex: cell elongation, apical dominance, general growth, and development) | Auxin efflux and polar auxin transport | STRING DB |
| **osj_34384** | Root hair | Root development | Probable GTP-binding protein | STRING DB |
| **pin1** | adventitious root | Regulation of adventitious root development via auxin pathway | Controls the traffic of auxin efflux carrier proteins | Literature |
| **pin2** | | Root growth angle | Component of the auxin efflux carrier | Literature |
| **pin3a** | Crown root | Root development, response to water stress | Auxin efflux and polar auxin transport | STRING DB |

| | | | | |
|---|---|---|---|---|
| **raa1** | Root system | Root development mediate by auxin | Cell cycle regulator during root development | STRING DB |
| **rcn1** | Lateral root | Promoting the outgrowth, hypodermal suberization of roots | Essential transporter for growth and development under abiotic stress, required for salt tolerance via Na/K homeostasis | STRING DB |
| **rhd3** | Root hair | Root development | Probable GTP-binding protein | Literature |
| **rr1** | Crown root | Root development | CK response regulators | Literature |
| **rr2** | Crown root | Root development | CK response regulators | Literature |
| **rr3** | Crown root | Root development | CK response regulators | Literature |
| **rr6** | Crown root | Suppressed root and vegetative development | CK response regulators | Literature |
| **sapk10** | Root hair | Produces longer root hairs | Aba activated protein kinase 10 production | Literature |
| **spl12** | Crown roots | Root development | Trans-acting factor | Literature |
| **spl3** | Crown roots | Increase number of roots | Trans-acting factor | Literature |
| **vln2** | | Root gravitropism | Modulation of polar auxin transport | Literature |
| **wox11** | Crown root | Root elongation, root development | CK-regulation | Literature |

**Supplementary Table B**:

List of NCBI gene symbols of proteins of extracted network module and the prediction scores for the top 75 candidates

| | **Protein** | **Gene symbol** | **Type of protein (seed/predicted)** | **Prediction score** |
|---|---|---|---|---|
| **1** | Os07t0108100-01 | LOC4342208 | Predicted | 398.2289137 |
| **2** | Osj_06916 | LOC4329481 | Predicted | 398.2289137 |
| **3** | Os05t0145700-01 | LOC4337795 | Predicted | 394.2489024 |
| **4** | Os05t0146100-01 | LOC4337797 | Predicted | 394.2489024 |
| **5** | Os05t0146900-00 | LOC9272414 | Predicted | 394.2489024 |
| **6** | GNP4 | LOC107277161 | Predicted | 278.8703058 |
| **7** | PDR16 | LOC4326812 | Predicted | 271.1123074 |
| **8** | LAX | LOC4327431 | Predicted | 262.3899959 |
| **9** | EXPA5 | LOC4330706 | Predicted | 198.1182047 |
| **10** | P0018C10.5 | LOC4325089 | Predicted | 198.1182047 |
| **11** | Os11t0540600-01 | LOC4350668 | Predicted | 198.1182047 |
| **12** | Os06t0610100-00 | Discontinued | Predicted | 198.1182047 |
| **13** | Os08t0423600-00 | LOC107276201 | Predicted | 198.1182047 |
| **14** | Osj_18177 | LOC 4338502 | Predicted | 196.1331962 |
| **15** | Osj_07614 | LOC4330055 | Predicted | 194.1581821 |
| **16** | Os11t0143300-02 | LOC4349747 | Predicted | 192.1931623 |
| **17** | HK6 | LOC107275680 | Predicted | 175.4249306 |
| **18** | C68 | LOC4328829 | Predicted | 174.1518585 |
| **19** | Os04t0672900-01 | LOC4337370 | Predicted | 174.1518585 |
| **20** | Os08t0467500-02 | LOC4345793 | Predicted | 174.1518585 |
| **21** | Os03t0253200-01 | LOC4332276 | Predicted | 174.1518585 |
| **22** | Os05t0107300-01 | LOC107275998 | Predicted | 173.900894 |
| **23** | P0425F05.2 | LOC4340552 | Predicted | 165.5780768 |
| **24** | Osj_34426 | LOC4350866 | Predicted | 157.7839569 |
| **25** | Os02t0190300-01 | LOC4328570 | Predicted | 157.7839569 |
| **26** | Os01t0967100-01 | LOC4324005 | Predicted | 156.1155519 |
| **27** | Osj_24793 | LOC4343642 | Predicted | 150.6678038 |
| **28** | HK4 | LOC4333916 | Predicted | 146.4983529 |
| **29** | Osj_09048 | LOC4331278 | Predicted | 144.1448717 |

| | | | | |
|---|---|---|---|---|
| **30** | HOX9 | LOC4348919 | Predicted | 141.028896 |
| **31** | Os09t0450600-00 | LOC4347222 | Predicted | 138.1439741 |
| **32** | Os01t0728200-01 | Discontinued | Predicted | 138.1439741 |
| **33** | Os07t0632600-01 | LOC4344004 | Predicted | 132.6048761 |
| **34** | Os07t0162600-02 | LOC4342462 | Predicted | 131.4163008 |
| **35** | Os03t0356470-00 | LOC9270588 | Predicted | 131.4163008 |
| **36** | Os03t0377300-01 | LOC9267254 | Predicted | 131.4163008 |
| **37** | Os03t0377500-01 | LOC9271339 | Predicted | 131.4163008 |
| **38** | Osj_34565 | LOC4350998 | Predicted | 131.4163008 |
| **39** | Os10t0485400-01 | Discontinued | Predicted | 131.4163008 |
| **40** | Os08t0547600-00 | LOC9269394 | Predicted | 131.4163008 |
| **41** | Osj_12688 | Discontinued | Predicted | 131.4163008 |
| **42** | Os08t0424100-00 | LOC9267362 | Predicted | 131.4163008 |
| **43** | P0025H07.5 | LOC4347247 | Predicted | 131.4163008 |
| **44** | EXPA31 | LOC4333168 | Predicted | 129.4387881 |
| **45** | Os11t0515500-01 | LOC4350590 | Predicted | 122.7141653 |
| **46** | PT2 | LOC4331637 | Predicted | 122.7141653 |
| **47** | Os01t0728150-00 | LOC107276471 | Predicted | 118.2806568 |
| **48** | PT3 | LOC4348740 | Predicted | 118.2806568 |
| **49** | Osj_32478 | LOC4349400 | Predicted | 115.2765157 |
| **50** | ARF19 | LOC4341978 | Predicted | 103.235112 |
| **51** | ORR5 | LOC4336439 | Predicted | 103.235112 |
| **52** | Os04t0445300-01 | LOC4335958 | Predicted | 102.7645228 |
| **53** | Os06t0697000-02 | LOC4341944 | Predicted | 102.7645228 |
| **54** | Os07t0582850-00 | LOC9266807 | Predicted | 98.06659812 |
| **55** | Os04t0101800-01 | LOC4334888 | Predicted | 98.06659812 |
| **56** | Osj_35591 | Discontinued | Predicted | 98.06659812 |
| **57** | Os09t0467700-01 | LOC4347323 | Predicted | 98.06659812 |
| **58** | CR4 | LOC4333525 | Predicted | 92.21651326 |
| **59** | Osj_22409 | LOC4341879 | Predicted | 90.21339716 |
| **60** | Os08t0564300-04 | LOC4346344 | Predicted | 89.18949213 |
| **61** | Os11t0498600-01 | LOC4350556 | Predicted | 88.88541299 |
| **62** | Os02t0235900-01 | LOC9271032 | Predicted | 86.45738679 |
| **63** | Osj_019622 | LOC9270361 | Predicted | 85.55435509 |
| **64** | Os11t0439600-01 | LOC4350420 | Predicted | 84.9841779 |

| | | | | |
|---|---|---|---|---|
| **65** | Os11t0672900-01 | LOC4351100 | Predicted | 84.9841779 |
| **66** | Os03t0303100-01 | LOC4332586 | Predicted | 84.9841779 |
| **67** | Osj_14626 | LOC4335696 | Predicted | 84.15088682 |
| **68** | Os01t0844300-01 | LOC4327507 | Predicted | 81.95702094 |
| **69** | Os05t0490600-00 | LOC4339167 | Predicted | 80.30566578 |
| **70** | OsGLHAT1 | LOC107275998 | Predicted | 80.30566578 |
| **71** | HOX10 | LOC4331345 | Predicted | 79.86774383 |
| **72** | IAA30 | LOC4352721 | Predicted | 79.86774383 |
| **73** | ASMT | LOC4346795 | Predicted | 78.05777595 |
| **74** | Osj_20964 | LOC4340753 | Predicted | 78.05777595 |
| **75** | Osj_30022 | LOC4347587 | Predicted | 78.05777595 |
| **76** | ADA2 | LOC4334126 | Seed | - |
| **77** | ARF12 | LOC4337363 | Seed | - |
| **78** | ARF25 | LOC4352783 | Seed | - |
| **79** | ARL1 | LOC9271993 | Seed | - |
| **80** | CCC1 | LOC4345272 | Seed | - |
| **81** | CINV1 | LOC4329626 | Seed | - |
| **82** | CKI1 | LOC4330018 | Seed | - |
| **83** | CKX4 | LOC4326515 | Seed | - |
| **84** | D14 | LOC4331983 | Seed | - |
| **85** | DRO1 | LOC4347169 | Seed | - |
| **86** | EL5.1 | LOC107276751 | Seed | - |
| **87** | EL5.2 | LOC4329685 | Seed | - |
| **88** | EXPA17 | LOC107276467 | Seed | - |
| **89** | EXPA8 | LOC4327217 | Seed | - |
| **90** | FH1 | LOC4325107 | Seed | - |
| **91** | GATB | LOC4350686 | Seed | - |
| **92** | GCN5 | LOC4348629 | Seed | - |
| **93** | GLR3.1 | LOC4336790 | Seed | - |
| **94** | GLU3 | LOC4336284 | Seed | - |
| **95** | GLU5 | LOC4324643 | Seed | - |
| **96** | GNA1 | LOC4347432 | Seed | - |
| **97** | IAA13 | LOC4334069 | Seed | - |
| **98** | IAA14 | LOC4334431 | Seed | - |
| **99** | MST3 | LOC4342198 | Seed | - |

| | | | | |
|---|---|---|---|---|
| **100** | MT2C | LOC4337596 | Seed | - |
| **101** | ORC3 | LOC4324643 | Seed | - |
| **102** | Os01t0682001-01 | LOC4324398 | Seed | - |
| **103** | Os12t0604600-01 | LOC4352733 | Seed | - |
| **104** | OsASL1 | LOC4332596 | Seed | - |
| **105** | OsEIL1 | LOC4332697 | Seed | - |
| **106** | Ost_28507 | LOC4346497 | Seed | - |
| **107** | Osj_34384 | LOC4350788 | Seed | - |
| **108** | PIN1 | LOC4330700 | Seed | - |
| **109** | PIN2 | LOC4341736 | Seed | - |
| **110** | PIN3A | LOC4326565 | Seed | - |
| **111** | RAA1 | LOC4326229 | Seed | - |
| **112** | RCN1 | LOC4332449 | Seed | - |
| **113** | RHD3 | LOC4326314 | Seed | - |
| **114** | RR1 | LOC4335937 | Seed | - |
| **115** | RR2 | LOC4329677 | Seed | - |
| **116** | RR3 | LOC4331245 | Seed | - |
| **117** | RR6 | LOC4337372 | Seed | - |
| **118** | SAPK10 | LOC4333435 | Seed | - |
| **119** | VLN2 | New submission | Seed | - |
| **120** | WOX11 | LOC4344325 | Seed | - |

**Supplementary Table C:** Details of missing seed proteins

| **Absent in raw dataset** | **Removed by filtering** |
|---|---|
| AIM1 | Osj_19760 |
| CARK | HO1 |
| SPL12 | SPL3 |

**Supplementary Table D:** Details of sub-modules

| SUB-MODULE | SEEDS | Functions of seeds | PREDICTED CANDIDATES |
|---|---|---|---|
| 1 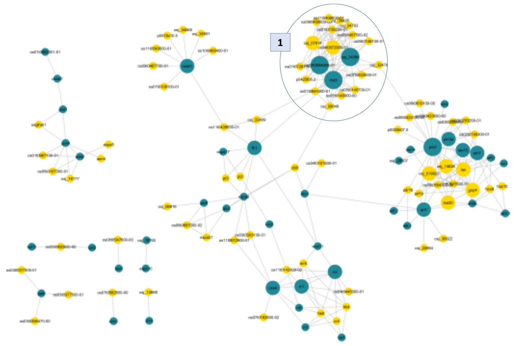 | RHD3 | Root hair development | Osj_32478 |
| | Osj_34384 | | Osj_07614 |
| | Os12t0604600-01 | | Osj_24793 |
| | | | P0425F05.2 |
| | | | Os01t0728200-01 |
| | | | Os07t0632600-01 |
| | | | Os05t0146900-00 |
| | | | Osj_09048 |
| | | | Os09t0450600-00 |
| | | | Os01t0728150-00 |
| | | | Os05t0145700-01 |
| | | | Os04t0672900-01 |
| | | | Os05t0146100-01 |
| | | | Os08t0467500-02 |
| | | | Osj_34384 |
| | | | Os01t0844300-01 |
| | | | Os11t0498600-01 |

| | | | | |
|---|---|---|---|---|
| **2** | 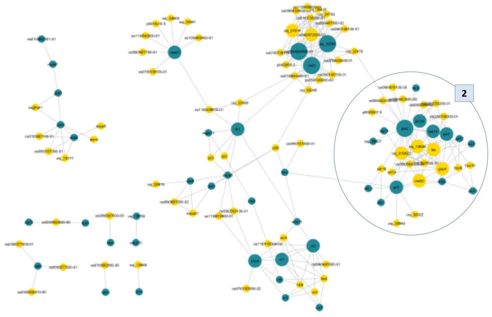 | EL5.2<br>EL5.1 | Degradation of cytotoxic proteins produced in root cells | ARF19<br>GNP4<br>HOX10<br>HOX9<br>IAA30<br>LAX<br>Os02t0190300-01<br>Os02t0235900-01<br>Os03t0253200-01<br>Os06t0610100-00<br>Os08t0423600-00<br>Os08t0424100-00<br>Os08t0564300-04<br>Os11t0515500-01<br>Osj_019622<br>Osj_14626<br>Osj_20964<br>Osj_30022<br>P0025H07.5<br>PDR16 |
| | | ARF25<br>ARF12 | Auxin response factors | |
| | | Osj_28507 | Influence in auxin-mediated developmental responses | |
| | | CCC1 | Osmotic regulation | |
| | | ARL1 | Auxin-mediated cell dedifferentiation | |
| | | PIN3A<br>PIN2<br>PIN1 | Auxin efflux and polar auxin transport | |
| | | IAA13<br>IAA14 | Members of the IAA family | |
| | | VLN2 | Modulation of polar auxin transport | |
| | | GLR3.1 | The glutamate-gate receptor may regulate cell proliferation and cell death in the root apex | |

| | | | |
|---|---|---|---|
| 3 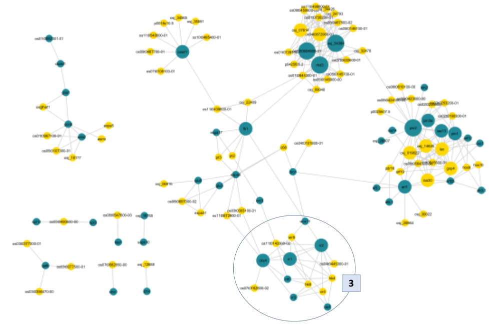 | CKI1 | Involved in ABA and brassinosteroid signaling pathways. | CR4<br>HK4<br>HK6<br>ORR5<br>Os04t0445300-01<br>Os07t0162600-02<br>Os11t0143300-02 |
| | RR6<br>RR1<br>RR2<br>RR3 | CK response regulators | |
| | CKX4 | CK signaling pathway | |
| | WOX11 | CK-regulation | |
| 4 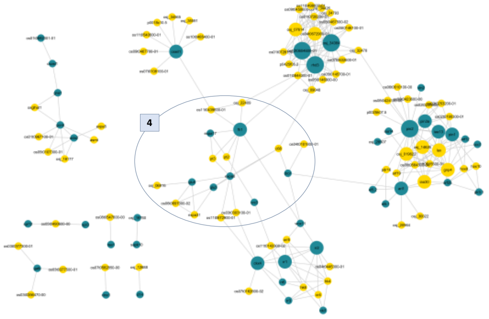 | FH1 | Root hair development | C68<br>EXPA31<br>Os03t0303100-01<br>Os04t0101800-01<br>Os06t0697000-02<br>Os11t0439600-01<br>Os11t0672900-01<br>Osj_06916<br>Osj_22409<br>PT2<br>PT3 |
| | GLU3 | Encoded for putative membrane-bound<br>Endo-1,4-β-glucanase, mutant lower the cellulose contents in its root cell walls | |
| | GLU5 | Endohydrolysis of (1->4)-beta-D-glucosidic linkages in cellulose | |
| | ORC3 | Component of the origin recognition complex that binds origins of replication | |
| | EXPA8 | Cell wall localized protein | |
| | EXPA17 | Cell wall remodeling | |
| | DRO1 | Enhances nitrogen uptake and cytokinin fluxes, regulating root growth angle | |

| | | | |
|---|---|---|---|
| **5** 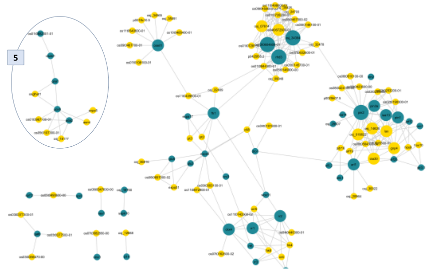 | GNA1 | Plays an important role in protein and lipid glycosylation | ASMT<br>EXPA5<br>Os01t0967100-01<br>Os05t0107300-01<br>OsGLHAT1<br>Osj_18177 |
| | ADA2 | Auxin or cytokinin signaling | |
| | Os01t0682001-01 | Reutilization of glutamine in developing organs. Plays a role in the development of tillers | |
| | gcn5 | Auxin and cytokinin signaling | |
| | osasl1 | Arginine biosynthesis | |
| **6** 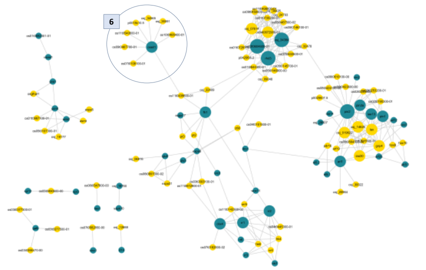 | oseil1 | Involved in the ethylene signaling pathway | Os07t0108100-01<br>Os09t0467700-01<br>Os10t0485400-01<br>Os11t0540600-01<br>OSj_34565<br>Osj_35591<br>P0018C10.5 |

*Figures*

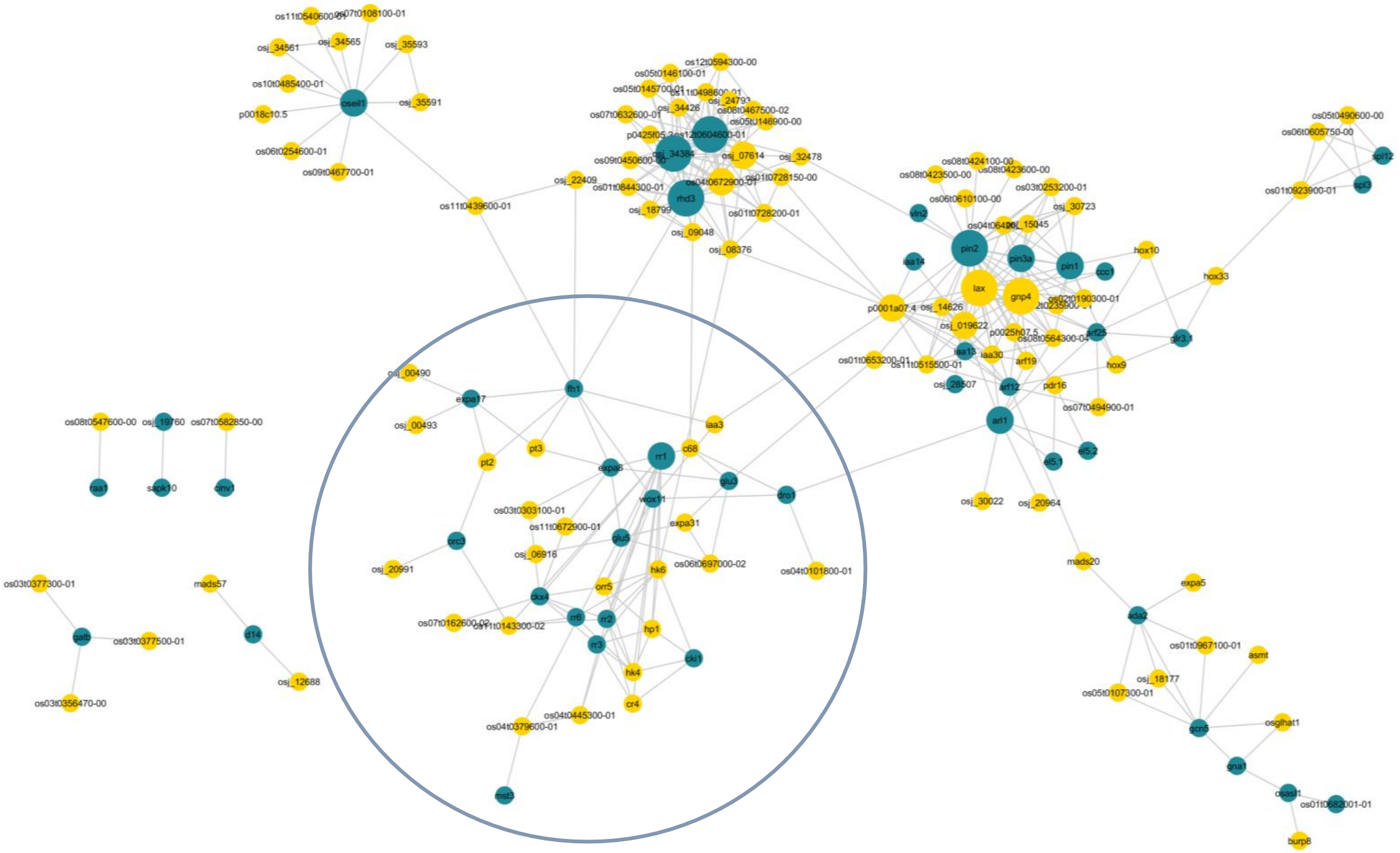


**Fig. A**. PPI network module visualization with 100 predicted proteins and 45 seed proteins. The Circled cluster indicates the integrated sub-module of sub-modules 3 and 4 in Fig.3. Seeds are represented by the cyan color nodes and predicted candidates are represented by the yellow color nodes
